# Cobalt induces the set-up of new structural networks in river biofilms: Impairment of autotrophic-heterotrophic coupling

**DOI:** 10.1101/2024.12.07.625411

**Authors:** Sarah Gourgues, Marisol Goñi-Urriza, Patrick Baldoni-Andrey, Nicholas Bagger Gurieff, Clémentine Gelber, Séverine Le Faucheur

## Abstract

Biofilms play a crucial role in biogeochemical cycles, making them essential to the functionality and stability of aquatic ecosystems. Their functioning is mainly driven by interactions between microorganisms that ensure the cycling of major elements. Evaluating the impact of stressors, such as metals, on these interactions is challenging. This study examined the effects of cobalt (Co) on the microeukaryotic community and their relationships with prokaryotes within biofilms grown in the presence of several Co concentrations (background concentrations, 0.1, 0.5, and 1 µM) for 28 days and 35 days after the end of Co injection. A previous work has suggested that *Cyanobacteria* were highly sensitive to Co leading to a reduction of the photosynthetic potential of the biofilm. In this study, the major primary producers, namely *Bacillariophyceae,* were also found to be sensitive to Co. The direct consequence of this sensitivity was an impairment of the autotroph-heterotroph coupling and the dominance of prokaryotic taxa in microbe-microbe interactions. The biofilms co-occurrence networks were then smaller, less connected but more centralized at 1 µM Co. Keystones, that were half affiliated with microalgae in the absence of Co, were mostly prokaryotes. As such, prokaryotes were found to control resource production and element cycling in Co-stressed biofilms. These changes in co-occurrence network organization and microbial community composition demonstrated a cascade of effects related to contamination in rivers. Our results further highlighted the strategy biofilms adopt to counteract the impairment of autotroph-heterotroph coupling, and to maintain their functions and ecosystemic roles in contaminated environments.

## 1. Introduction

Freshwater biofilms are highly diverse microbial communities including algae, bacteria, fungi, and meiofauna embedded in a matrix of extracellular polymeric substance (EPS), which develop on submerged substrata (1,2). Biofilms are at the bottom of the trophic chain and contribute to energy flows, nutrient cycles, and global biogeochemical cycles of major (C, N, P, S) and trace elements in fluvial ecosystems (3,4). These biological systems are critical in organic matter cycling due to the close interactions between autotrophs and heterotrophs in the biofilm. The autotrophic component comprises mainly of photosynthetic organisms such as green algae and cyanobacteria, which perform photosynthesis (5). The produced organic matter is then mineralized/transformed by heterotrophic organisms such as bacteria and fungi (6). In turn, the CO2 produced during the transformation of organic matter by heterotrophs is available for phototrophs. As a highly metabolic active system, biofilms are also strongly involved in other nutrient cycling to respond to the biotic demand of microorganisms (4). Biofilm structures are defined by the interactions between microorganisms, that enable them to function and provide ecosystem services. However, most of the studies investigating the contaminant effect on biofilms focus on microbial composition characterization of prokaryotes or microalgae, without considering that biofilms can be regarded as a unique biological consortium that interact and operates like a meta-organism (7).

The modification of the community structure and function that is usually described following stress can be a direct consequence of stressors and/or an indirect consequence of the microorganisms’ interactions/network. Several studies have investigated the effect of environmental changes on microbial co-occurrence in soils (8–10) and sediments (11–13) and showed that the structure of microbial networks (including network size, complexity, or keystone number) was affected by stresses such as algal bloom, water contamination by hydrocarbons, or agricultural intensification. Although several studies have assessed the response of microbial communities’ structure and functions in freshwater biofilms under stressed conditions (14,15), consideration of microbial interactions and analysis of co-occurrence networks has not been adequately addressed. The study of microbial interactions is essential to provide a complete picture of ecological repercussions induced by stresses on biofilms.

Cobalt (Co) is an emerging contaminant widely used in the context of the energy transition. The increasing production of lithium-ion batteries has led to an exceptional rise in its extraction rates and use in the last decades (16–18). Its increasing detection in aquatic systems at concerning concentrations has led to its classification as a toxic substance in Canada with the implementation of Federal Water Quality Guidelines to protect aquatic life (19). It has also been identified as a relevant substance to monitor in continental waters in France (20). To date, most of the studies investigating Co impacts on microbial life were performed on prokaryotic and microalgal model organisms (21,22) and only a few investigated Co impacts on biofilms. Cobalt bioaccumulation in biofilms has been reported to be linked to a disruption of chlorophyll a synthesis and biofilm growth at concentrations of 85 and 17 µM Co, respectively (23). Another study demonstrated that the composition of the prokaryotic and micro-eukaryotic communities and the metabolomic response were affected rapidly (within hours) in mature biofilms translocated in 0.5 and 1 µM Co-contaminated waters (24).

Recently, we grown biofilms in artificial streams contaminated with different Co concentrations and investigated the impact of Co on the prokaryotic members within biofilms. We reported that Co contamination in freshwaters at environmentally relevant concentrations (0.5 and 1 µM Co) strongly impacted the structure and functional potential of the prokaryotic component of biofilms (25). In this previous study, we demonstrated that *Cyanobacteria* were highly sensitive to Co, and consequently, an important reduction of the photosynthetic potential of the biofilm was suggested. At the same time, the prokaryotes potentially involved in carbon fixation using alternative pathways to photosynthesis increased. Although this work provided insights into the impacts of Co on freshwater biofilms, it considers prokaryotes alone and gives only a narrow picture of the complex interplays between microbial components. Indeed, the microbial communities within biofilms are strongly associated, and microalgae are a major component involved in carbon fixation by photosynthesis. It is therefore crucial to consider microeukaryotic component alongside prokaryotes from the same biofilm, particularly their interactions, to gain a more complete understanding of the impact that Co can have on river biofilms and their ecological roles.

The present study aimed to investigate the Co effects on the (i) structure of the microeukaryotic community developing within river biofilms, and (ii) microbial interactions occurring at the biofilm level focusing on the autotroph-heterotroph coupling. For this purpose, previous biofilms grown in artificial streams enriched with Co at environmentally relevant concentrations were examined for the abundance and structure of their microeukaryotic community at 7, 14, 21 and 28 days of growing and after 35 days of recovery without any Co addition. Then, the relationships between prokaryotic (25) and microeukaryotic communities were examined using co-occurrence network properties to evaluate how their interactions could be affected. We demonstrated that Co induced a significant reorganization of the co-occurrence network, leading to the alteration of the essential autotroph-heterotroph coupling. The sensitivity of the main primary producers from exposure to Co contamination led to the prevalence of prokaryotes as the main drivers of microbial interactions in phototrophic biofilms.

## 2. Material and methods

### 2.1. Biofilm growth and sampling

The experiment was carried out in outdoor artificial streams (Pilot Rivers) at TotalEnergies facilities in Lacq (PERL, France) (Figure S1) (26). Water from the Gave de Pau River continuously fed the artificial streams. This water was characterized by a decreasing temperature (from 14.3 ± 0.1 °C to 8.17 ± 0.0 °C from the start to the end of the experiment, respectively), an average pH of 8.3 ± 0.3, and an average background total dissolved Co concentration of 4.62 ± 4.02 nM for 28 days. Its average concentration of dissolved organic carbon was 0.39 ± 0.19 mg.L^−1^ whereas the average concentrations of Ca^2+^ and Mg^2+^ were 8.98 ± 0.03 × 10^−4^ M and 1.6 ± 0.1 ×10^−4^ M, respectively. All physico-chemical parameters measured during the experiment are given in (25). A random selection of 12 streams (40 m long, 0.5 m wide, 0.5 m deep, and with a flow rate set at 7.5 m^3^.h^−1^) was made to assign a Co exposure concentration. Solutions of cobalt (Co(II) chloride hexahydrate, 98%, Thermo Scientific Chemicals) was continuously injected into nine streams to achieve nominal concentrations of 0.1, 0.5 and 1 µM in triplicate (3 treatments x 3 streams). The injection of Co started one day before the exposure experiment started and continued for a period of 29 days. Three additional streams with only background Co concentrations from the Gave de Pau River were used as controls. The exposure concentrations remained stable over the experiment (25). Thirteen sterile glass slides (10 x 20 cm) were placed into each stream at the beginning of Co exposure to serve as a substrate for biofilm growth. Biofilms and water were then collected every 7 days during biofilm growth (D7, D14, D21, and D28) as well as 35 days after the Co injection was stopped (DR63). Subsamples from collected biofilms were used to characterize the microeukaryotic community of river biofilms. Samples were weighed and stored at −80°C for molecular analyses. Prokaryotic community characterization, dry weight and Co bioaccumulation in biofilms were published elsewhere (25). Briefly, the dry wet biomass increased over the 28 days of exposure regardless of Co concentrations. The bioaccumulated levels of Co were linearly correlated with ambient Co concentrations and ranged from 0.09 ± 0.01 to 9.25 ± 1.19 µmol Co.g^−1^ DW for background Co concentration and 1 µM Co, respectively.

### 2.2. Diversity of microeukaryotic communities

Extraction of DNA from biofilms was performed using PowerSoil DNA Isolation Kits (Qiagen, Germany) following the manufacturer’s instructions. Quantification of extracted DNA was done using Qubit 1X dsDNA BR assay kits (ThermoFisher Scientific, USA). Hypervariable regions (V8-V9) of 18S rRNA gene were then amplified by PCR using primers V8-1422F and V9-1797R (27) and the following protocol: initial denaturation of 10 min at 95°C, followed by 25 cycles of 15 s denaturation at 95°C, 30 s annealing at 57°C and 30 s of elongation at 72°C. A final elongation of 7 min at 72°C was performed. Sequencing of amplicons was performed by *La Plateforme Génome Transcriptome de Bordeaux* (*PGTB*, Bordeaux, France) using the Illumina Miseq 2 x 300 bp method and reagent kit V3. Raw sequences were submitted to the National Center for Biotechnology Information as a Sequence Read Archive (SRA) under the accession number PRJNA1166897.

Processing of the raw sequences was performed using the FROGS (version 4.1.0) pipeline (Find Rapidly OTUs with Galaxy Solution) (28). Steps of merging, denoising and dereplication of raw sequences were performed using the VSEARCH (29) tool and sequences with less than 200 bp were removed for downstream analyses. Clustering in operational taxonomic units (OTUs) was done with SWARM (30), setting the aggregation distance clustering to 1. Chimeras and OTUs representing less than 0.005% of all sequences were removed (31). Data was normalized to 10,789 reads per sample and the final total number of OTUs was 1,178. The reference database 18S SILVA 138.1 was used for the taxonomic assignments of OTUs.

### 2.3. Quantification of ribosomal 18S gene

Quantification of 18S rRNA gene copy number was used as a proxy for the abundance of microeukaryotes. Real time PCR was performed with DyNAmo Flash SYBR Green qPCR Kits (ThermoFisher Scientific, USA) with the same primers as those used for microeukaryotic diversity analysis without the Miseq adaptors. RT-PCR was performed in a final volume of 20 µL using 0.8 µM of each primer and 1 µL of DNA template (between 7.2 and 412 ng). Amplification was done as follows: initial denaturation of 10 min at 95°C followed by 25 cycles of 15 s of denaturation at 95°C, 30 s of annealing at 57°C and 30 s of elongation at 72°C. Amplification specificity was checked, producing melting curves. Absolute quantification was performed with home-made standards consisting of pGEM cloned amplicons of *Chlamydomonas reinhardtii* 18S rRNA gene. Analyses were performed in triplicates for each biological replicate. The efficacity of the quantitative PCRs was of 93 %.

### 2.4. Construction of co-occurrence networks

Co-occurrence relationships between microorganisms growing within biofilms were assessed at the different Co exposure concentrations. Biofilms sampled during the first 28 days of growth (four sampling dates: 7, 14, 21 and 28 days of Co exposure) were grouped according to the Co exposure concentrations *i.e.* control conditions (4.62 ± 4.02 nM Co, n=34), 0.1 µM Co (n=35), 0.5 µM Co (n=36) and 1 µM Co (n =36). Co-occurrence networks are based on correlation coefficients calculated between pairs of organisms, representing positive and negative relationships between both organisms. Spearman’s correlations were determined between pairs of OTUs with the *corTest* function from the *psych* package in Rstudio and corrected using the Benjamini-Hochberg false discovery rate (FDR) correction (32). Only statistically (adjusted p < 0.05) and robust correlations (Spearman’s correlation coefficient; ρ > |0.8|) were selected. Cytoscape v.3.10.2 was then used for network visualization and topological features calculation with the *analyse network* tool (33). To assess whether the four constructed networks (one for each Co exposure condition) were not random and represented true microbial interactions, one thousand additional random networks were generated for each network using the Erdös-Renyi null-model |*G (n,m)*| with the same number of nodes (n) and edges (m) as the initial studied network, with the *erdos.renyi.game* function from *igraph* package in Rstudio. The topology of resulting networks was also evaluated with the calculation of topological parameter nodes degree, with the *igraph* package of Rstudio. The detection of keystone taxa was carried out as recommended with the selection of OTUs (nodes) with a high degree and closeness and low betweenness centrality, using a threshold of the third percentile of each parameter score (34,35).

### 2.5. Statistical analyses

Alpha diversity was assessed by calculating species richness, diversity (Shannon index), and evenness (Pielou index) using the Vegan package in RStudio (V2023.12.1.) (36). ANOVA and Tukey post-hoc tests were applied in RStudio for all comparisons, when applicable. If conditions for application were not met, the non-parametric Kruskal-Wallis and subsequent Dunn post-hoc tests were applied. Non-metric multi-dimensional scaling (NMDS) based on Bray-Curtis dissimilarity distances between OTUs was used to assess beta-diversity. Permutational multivariate analysis of variance (PERMANOVA) from the Vegan package was applied to assess significant differences between samples and followed by pairwise comparisons using pairwiseadonis2 package (37). The difference of main taxa (representing > 1% of the community) related to Co exposure were analyzed using the statistical analysis of the metagenomic profiles (STAMP) tool, using White’s non-parametric test and an effect size filter for proportion differences exceeding 1% (38). To evaluate the effect of dominant phototrophic groups on the prokaryotic community structure, a canonical correspondence analysis (CCA) was performed using the Vegan package (36). Phototrophic groups (*cyanobacteria* and microalgae) and Co concentration were treated as explanatory variables while the prokaryotic OTUs served as response matrix. Multicollinearity among variables was assessed prior analysis using the Variance Inflation Factor (VIF) function with a threshold of 10. The statistical significance of the model and individual variables were tested using permutation-based ANOVA with 999 permutations. The distribution of topological parameters from empirical networks was compared with those randomly generated using the Kolmogorov-Smirnov test from *stats* package (39) to determine whether the networks produced in the study were not random.

## 3. Result

### 3.1. Structure of the microeukaryotic community in biofilms growing at different Co concentrations

#### 3.1.1. Abundance of the microeukaryotic community

The copy number of the 18S rRNA was used as a proxy of microeukaryotic abundance (Figure 1, Table A1). In control biofilms, the microeukaryotic abundance increased significantly between D7 and D14 (p < 0.05) reaching 4.76 × 10^6^ ± 2.71 × 10^6^ 18S copies.cm^−2^. After D14, the 18S rRNA copy numbers slowly decreased with a significant decrease between D21 and D28 as well as after the recovery period (p < 0.05). Microeukaryotes were less abundant at D7 in biofilms exposed to 0.5 and 1 µM Co (p < 0.05), and at D21 in biofilms exposed to 0.1, 0.5 and 1 µM Co (p < 0.05). After the end of Co exposure, the microeukaryotic abundance declined in all biofilms and no difference in abundance was measured when comparing previous Co exposure conditions.

**Figure 1:**
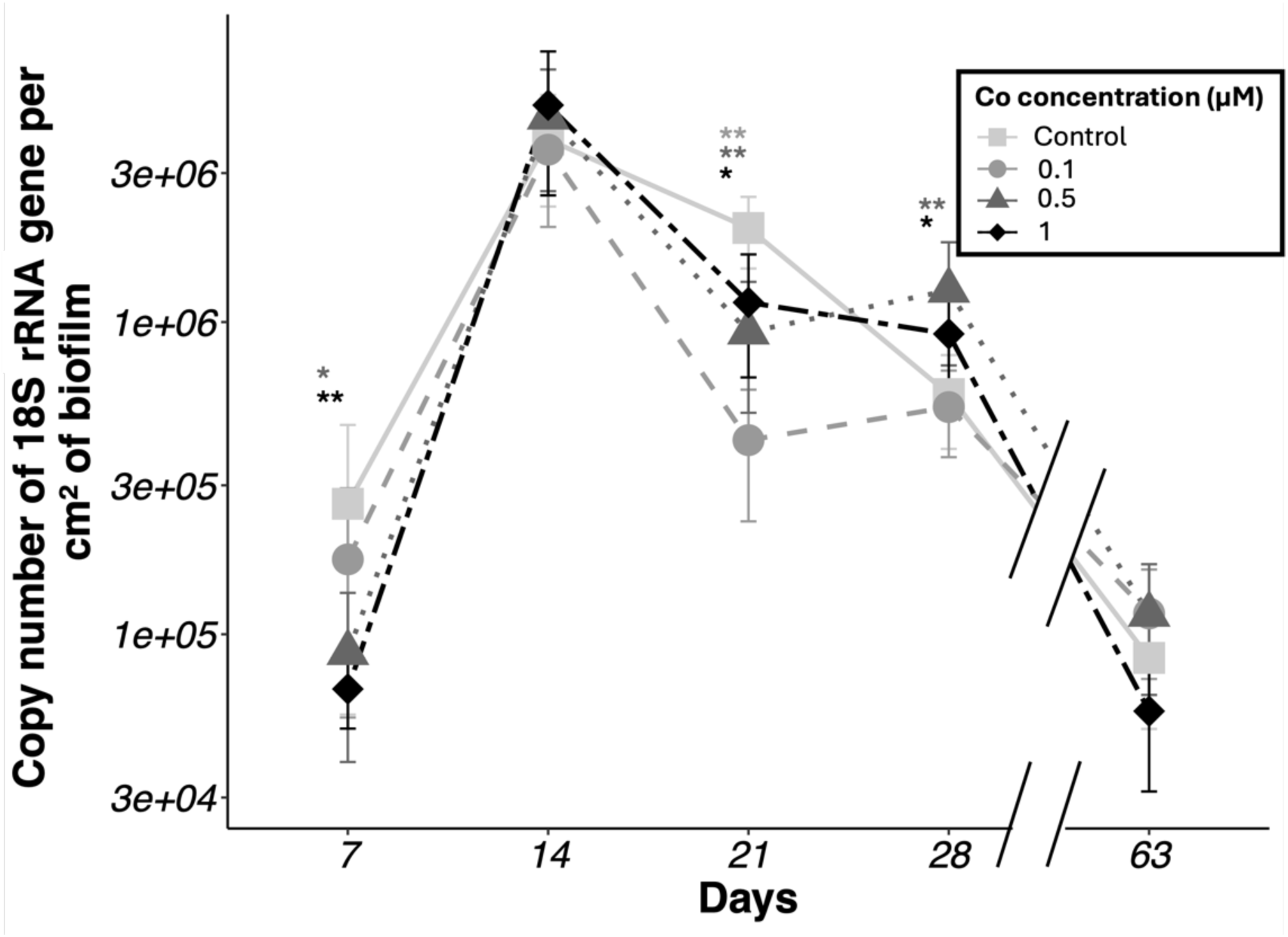
18S rRNA gene copy number in biofilms grown in several concentrations of of Co during 28 days and during 35 additional days after the Co injection was stopped (Day 63, DR63). The shape of dots and lines indicate the increasing Co concentrations by comparison with control conditions (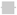: Control conditions; 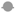: 0.1 µM Co; 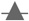: 0.5 µM Co; 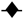: 1 µM Co).

#### 3.1.2. Diversity of microeukaryotic communities

Microeukaryotic community was dominated by microalgae and meiofauna, which represented 56 ± 13 % and 15 ± 9 % of the community respectively, regardless of sampling time or exposure conditions (Table A2). Fungi and macroalgae were also found but represented less than 5% of the eukaryotic community. Microalgae and meiofauna communities were selected for deeper analysis due to their predominance in biofilms and because they are representative of microorganisms belonging to different trophic levels.

The specific richness of the microalgal communities significantly increased between D7 and D14 (p < 0.05) regardless of Co concentrations before remaining stable or slightly decreasing until D28 (Table A3). Biofilms growing in the presence of 0.5 µM Co had significantly higher specific richness, Shannon and evenness (Pielou) indexes at D21 and D28 (p < 0.05) than the control biofilms. After 35 days of stopping Co exposure, microalgal community was characterized by a lower specific richness and diversity indexes (p < 0.05) compared with biofilms at D28. However, they remained higher in biofilms previously exposed to 0.5 and 1 µM Co. For the meiofaunal community (Table A4), diversity metrics in control biofilms were relatively stable when compared to biofilms exposed to Co, with the highest values of specific richness and diversity indexes at D14. In the presence of Co, meiofaunal richness and diversity indexes declined from D14 to D28 (p < 0.05). The end of the Co exposure led to a significant decrease of diversity metrics in all conditions (p < 0.05) without being able to determine a clear long-term effect of Co exposure.

Beta-diversity of microalgal and meiofaunal communities (based on Bray-Curtis distances) are represented on the two-dimensional Non-Metric Multi-Dimensional Scaling (NMDS) ordination (Figure 2). Samples were clustered according to exposure time (axis NMDS 1) and Co exposure concentration (axis NMDS 2). Both communities generally had changing structures during biofilm growth, regardless of Co concentration (p < 0.05) (Table A5-A6), with a few exceptions between D21 and D28. Cobalt exposure had also a significant effect (p<0.05) on community structure. While biofilms growing in control and 0.1 µM Co had similar microalgal and meiofauna communities, biofilms exposed to 0.5 µM and 1 µM had different community compositions in comparison with the control (p < 0.05) (Table A5-A6). The differences between these clusters increased with time, as observed in the NMDS ordinations by the increasing distances (Figure 2). These structural differences remained after the 35 days of recovery (DR63) after stopping Co dosing (Figure 2) (Table A5-A6; p < 0.05).

**Figure 2:**
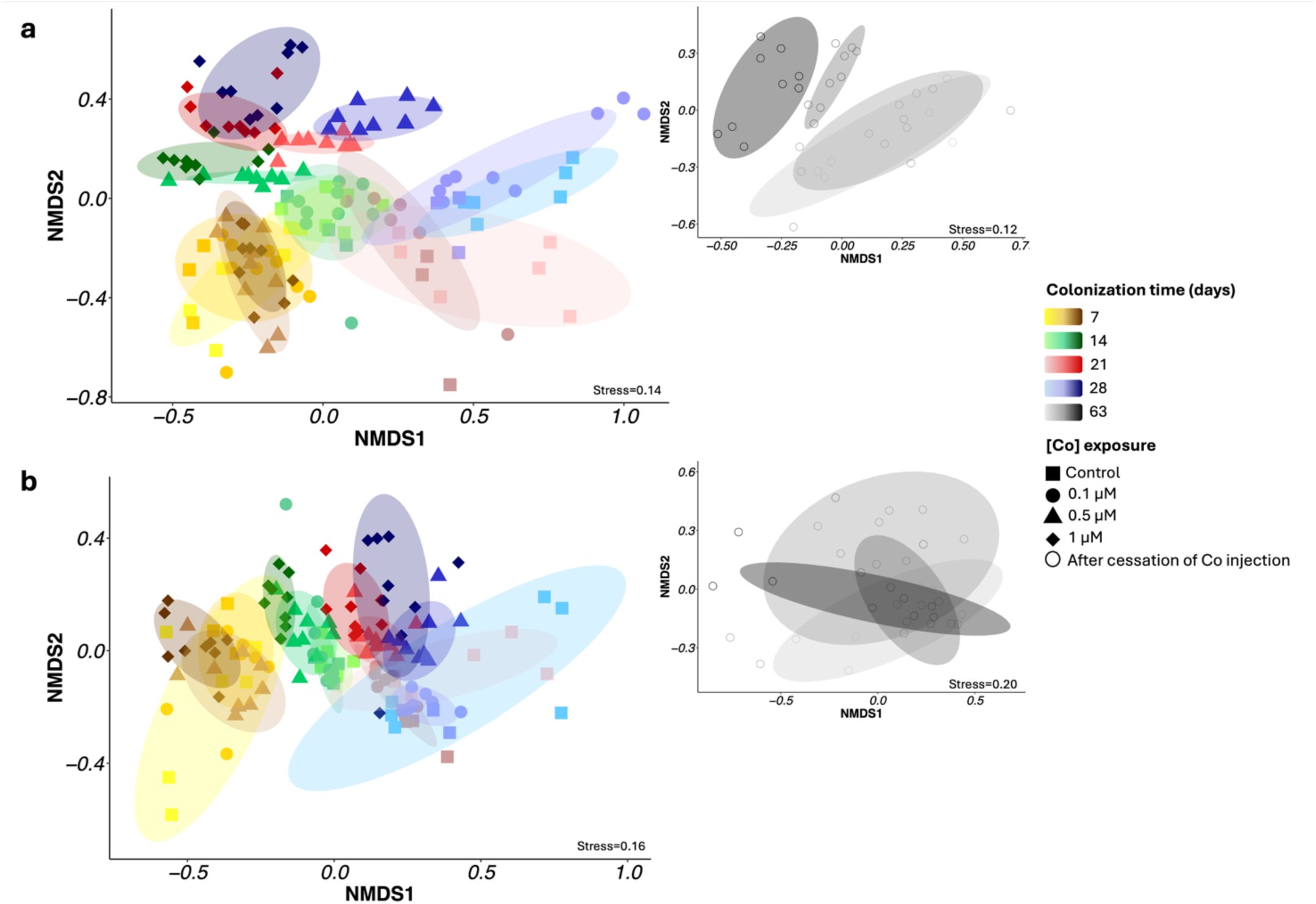
Non-metric multi-dimensional scaling ordination based on Bray-Curtis dissimilarity matrix of OTUs from microalgal community (stress= 0.12) (**a**) and meiofaunal community (stress=0.20) (**b)**. Sampling times are represented as follows: Day 7 (yellow), Day 14 (green), Day 21 (red), Day 28 (blue), and DR63 after cessation of Co injection (black). Conditions of Co exposure are represented as follows: ∎ Control (background concentration); ● 0.1 µM; ▲0.5 µM; ♦ 1µM. Ellipses include 80% of the samples from the same Co condition exposure and sampling time

The microalgal community of control biofilms was mainly dominated by *Bacillariophyceae, Coscinodiscophyceae, Chlorophyceae,* and *Cryptophyceae* (Figure S2a). The meiofauna was dominated by *Intramacronucleata, Monogononta, Ostracoda* and *Thecofilosea* (Figure S2b). The composition of biofilms exposed to 0.1 µM Co was not significantly different from the control. In contrast, those exposed to 0.5 µM and 1 µM Co highlighted the resistance of the *Cryptophyceae* during biofilm growth and of the *Chlorophyceae* at later stages (D21 and D28) (Figure S3a, b). A sensitivity of the *Bacillariophyceae* and the *Coscinodiscophyceae* was also observed (Figure S3c, d). Significant resistance trends were observed in the meiofauna over biofilm growth in the presence of 0.5 µM and 1 µM Co for the *Intramacronucleata* (D14, D21)*, Glissomonadida* (D21 and D28), and *Thecofilosea* (D28) (Figure S4). After 35 days of recovery (DR63), biofilms grown at 1 µM Co remained different with less *Bacillariophyceae* while the relative abundance of *Chlorophyceae, Ulvophyceae* and *Intramacronucleata (*meiofauna) increased.

### 3.2. Cobalt impacts on co-occurrence networks in biofilms

#### 3.2.1. Cobalt affects topological features and composition of co-occurrence networks

Co-occurrence networks were constructed for each condition of Co exposure (Figure 3) (i.e. all samples collected between D7 and D28 combined for each Co concentration) and their topological features are summarized in Table 1. The network constructed for control biofilms had 497 nodes and 2,662 edges. A lower proportion of the community was involved in the network with increasing Co concentrations. Reduction of the overall network connectivity was also described by a decrease of the network diameter and of the average number of neighbours per node. Nevertheless, the connectance between nodes increased with Co concentration. Indeed, the network was denser at 1 µM Co (0.04) than in control conditions (0.02), and tended towards the formation of sub-networks where nodes interact more, as indicated by the clustering coefficient. The network heterogeneity, which measures the variability in connectivity among nodes, decreased from 1.40 in the control to 1.13 at 1 µM Co, and network centralization around a few central nodes was also enhanced in biofilms exposed to Co compared with the control biofilm (Table 1).

**Figure 3:**
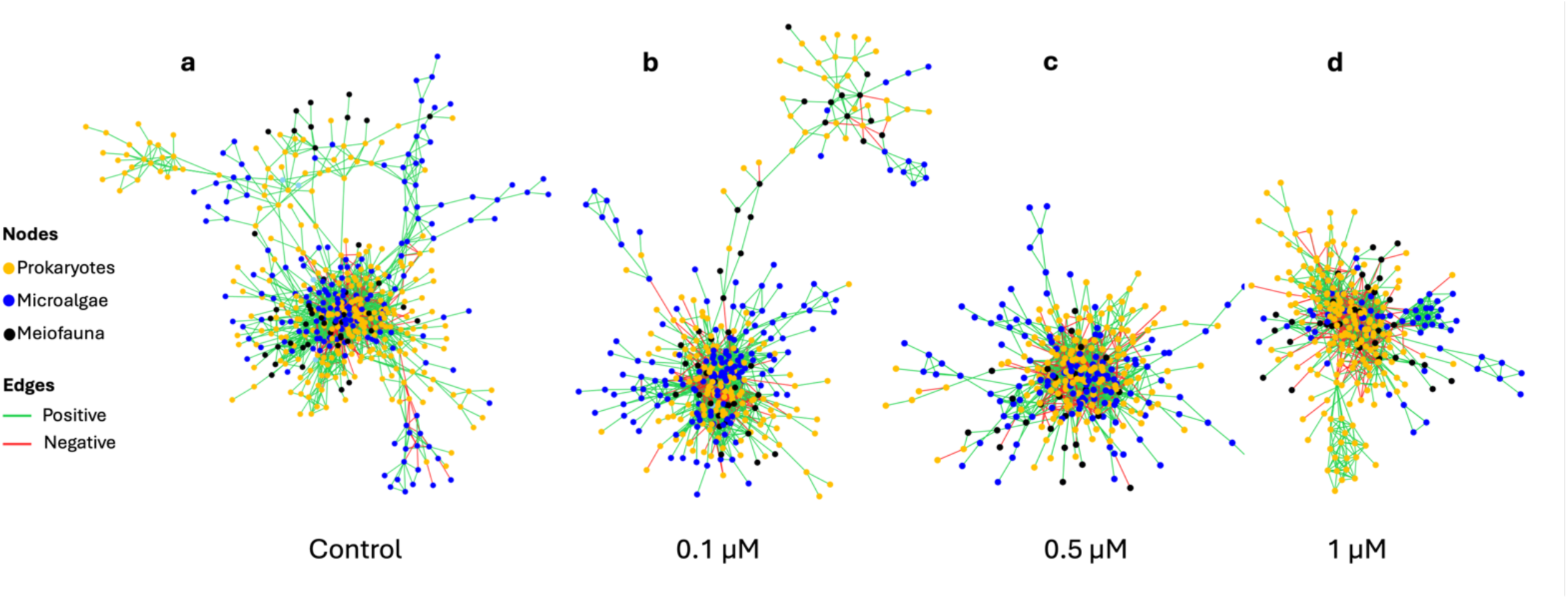
Co-occurrence networks between prokaryotic (yellow), microalgal (blue) and meiofaunal (black) taxa of biofilms grown under different Co concentrations: Control conditions (**a**), 0.1 µM Co (**b**), 0.5 µM Co (**c**) and 1 µM Co (**d**). Associations between taxa were generated with Spearman’s correlations. Strong (ρ > | 0.8 |) and significant (p < 0.05) correlations were selected for network construction in each condition. Green lines: positive correlations; red lines: negative correlations

**Table 1:** Topological features of co-occurrence networks constructed from microbial communities of biofilms exposed to different Co concentrations: Control (n=34), 0.1 µM (n=35), 0.5 µM (n=36) and 1 µM Co (n=36). One network includes all the sampling times during exposure (from D7 to D28) for a given concentration. Spearman’s correlations were calculated between pairs of OTUs and only significant (p < 0.05) and robust correlations ρ > |0.8| were selected for networks construction and calculation of topological parameters.

| Topological features | Control | 0.1 $\mu\text{M}$ | 0.5 $\mu\text{M}$ | 1 $\mu\text{M}$ |
| --- | --- | --- | --- | --- |
| Proportion of community involved (%) | 65% | 56% | 52% | 46% |
| Connected component | 1 | 1 | 1 | 1 |
| Number of nodes | 497 | 362 | 300 | 279 |
| Number of edges | 2,662 | 2,281 | 1,518 | 1,377 |
| Positive edges | 2,026<br>(76%) | 1,689<br>(74%) | 1,014<br>(67%) | 909<br>(66%) |
| Negative edges | 636<br>(24%) | 592<br>(26%) | 504<br>(33%) | 468<br>(34%) |
| Average number of neighbours | 10.71 | 12.60 | 10.12 | 9.87 |
| Network diameter | 16 | 17 | 11 | 9 |
| Network radius | 9 | 9 | 6 | 5 |
| Characteristic path length | 4.85 | 5.32 | 3.54 | 3.60 |
| Clustering coefficient | 0.37 | 0.40 | 0.37 | 0.43 |
| Network density | 0.02 | 0.04 | 0.3 | 0.4 |
| Network heterogeneity | 1.40 | 1.36 | 1.27 | 1.13 |
| Network centralization | 0.15 | 0.20 | 0.23 | 0.19 |
| Number of keystone taxa | 11 | 12 | 2 | 4 |

Microbial community composition within networks also varied as a function of Co exposure (Figure S5). Control networks were dominated by prokaryotic nodes (56%) followed by microalgae (35%) and meiofauna (9%). The most significant shift was observed in networks built at 1 µM Co where 67% of the nodes were prokaryotes and only 21% were microalgae. The proportion of nodes associated with meiofauna ranged from 8% to 12%.

#### 3.2.2. Identification and dynamics of keystone taxa

Nodes with high degree, high closeness centrality and low betweenness centrality were defined as keystones (34). A decreasing number of keystones were found with increasing Co concentrations, from 11 keystones in control biofilms to 4 at 1 µM Co (Table 1,2). Keystone taxa were almost half microalgae and half prokaryotes in the control, 0.1 µM Co and 0.5 µM Co networks whereas they were mainly composed of prokaryotic OTUs at 1 µM. The keystone dynamics, taxonomy, and modification in their relative abundance are provided in Table 2.

**Table 2:**
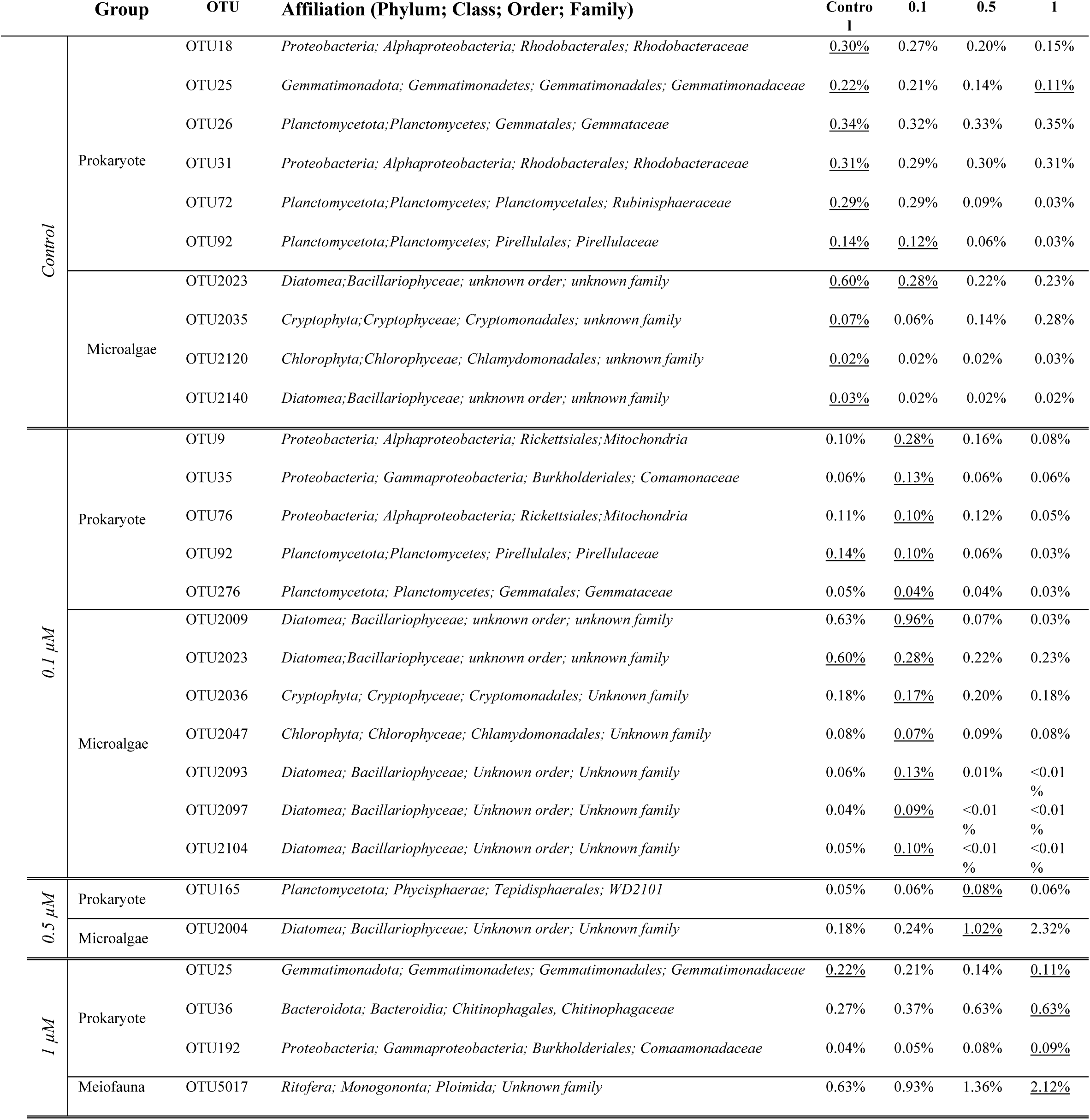
Keystone taxa in co-occurrence networks constructed from biofilms growing at each condition of Co exposure (Control, 0.1 µM Co, 0.5 µM Co, 1 µM Co). Their taxonomical affiliation and relative abundance in the community as a function of Co concentration are presented. The underlined relative abundance value indicates that the OTU is keystone at the related Co concentration condition.

The topological parameters (degree and betweenness centrality (BC)) dynamics of the keystone taxa identified in the control network were examined as a function of increasing Co concentrations (Figure S6a, b). Three types of responses were identified : Group 1 including 5 OTUs (18, 72, 92, 2009 and 2035), that disappeared from the network composition at 0.5 µM or 1 µM; Group 2 including 4 control keystone OTUs (31, 2023, 2120 and 2140) that were involved in almost all of the networks, but had decreasing number of neighbours and BC with increasing Co concentrations; and, Group 3 including 2 OTUs (25 and 26) that showed a U-bell type of response with a decrease of degree and BC at 0.1 µM or 0.5 µM Co. The relative abundance of all the control keystone OTUs was investigated within the overall microbial community of each studied biofilm (Figure S6c; Table A7). The relative abundance of 6 keystones OTUs (18, 25, 72, 92, 2009 and 2023) decreased in biofilms when Co concentrations increased. In contrast, the relative abundance of the OTU2035 increased and 4 OTUs (26, 31, 2120 and 2140) had a stable relative abundance across all of the exposure conditions. Among the 11 keystones of the control network, three (OTU92, OTU2009, OTU2023) were also keystones in the network at 0.1 µM while one (OTU25) was keystone in both control conditions and at the highest Co exposure (1 µM). This bacterial OTU was affiliated with the *Gemmatimonadales* from the *Gemmatimonadota* phylum.

The analysis of the OTU25 sub-network highlighted 66 and 59 direct neighbours in control and 1 µM Co conditions, respectively (Figure S7; Table S1); most of these neighbours were specific to the Co exposure concentration (Figure S7). Overall, a decrease of the interactions between OTU25 and microalgal OTUs at 1 µM was found, and the still-present interactions with *Bacillariophyceae* switched from positive to negative (Table S1). For prokaryotic neighbours of OTU25, while interactions with members of the *Planctomycetes* decreased with increasing Co concentrations, the number of interactions with *Bacteroidia* (*Sphingobacteriales* and *Flavobacteriales* principally) highly increased at 1 µM Co and these interactions switched from negative in the control condition to positive in 1µM of Co condition (Table S1).

#### 3.2.3. Microbial interactions in response to Co contamination in river biofilms

The interactions between prokaryotes, microalgae, and meiofauna (≥ 1% of the overall interactions) are represented for each exposure condition in a heatmap (Figure 4; Table A8-A11). In control conditions (Figure 4a; Table A8), most of the interactions involved *Bacillariophyceae*, *Alphaproteobacteria,* and *Planctomycetes* accounting for 26%, 25% and 21% of total interactions, respectively. For biofilms growing in the presence of 0.1 µM and 0.5 µM Co, the proportions of interactions involving *Bacillariophyceae* (diatoms) increased up to 50% of the interactions (Figure 4b, c; Table A9, A10), but this proportion collapsed to 15% within biofilms exposed to 1 µM of Co (Figure 4d; Table A11). Conversely, *Bacteroidia* members became the most dominant group in microbial interactions at 1 µM, being involved in almost 40% of interactions while this interaction represented only 17% in the control network.

**Figure 4:**
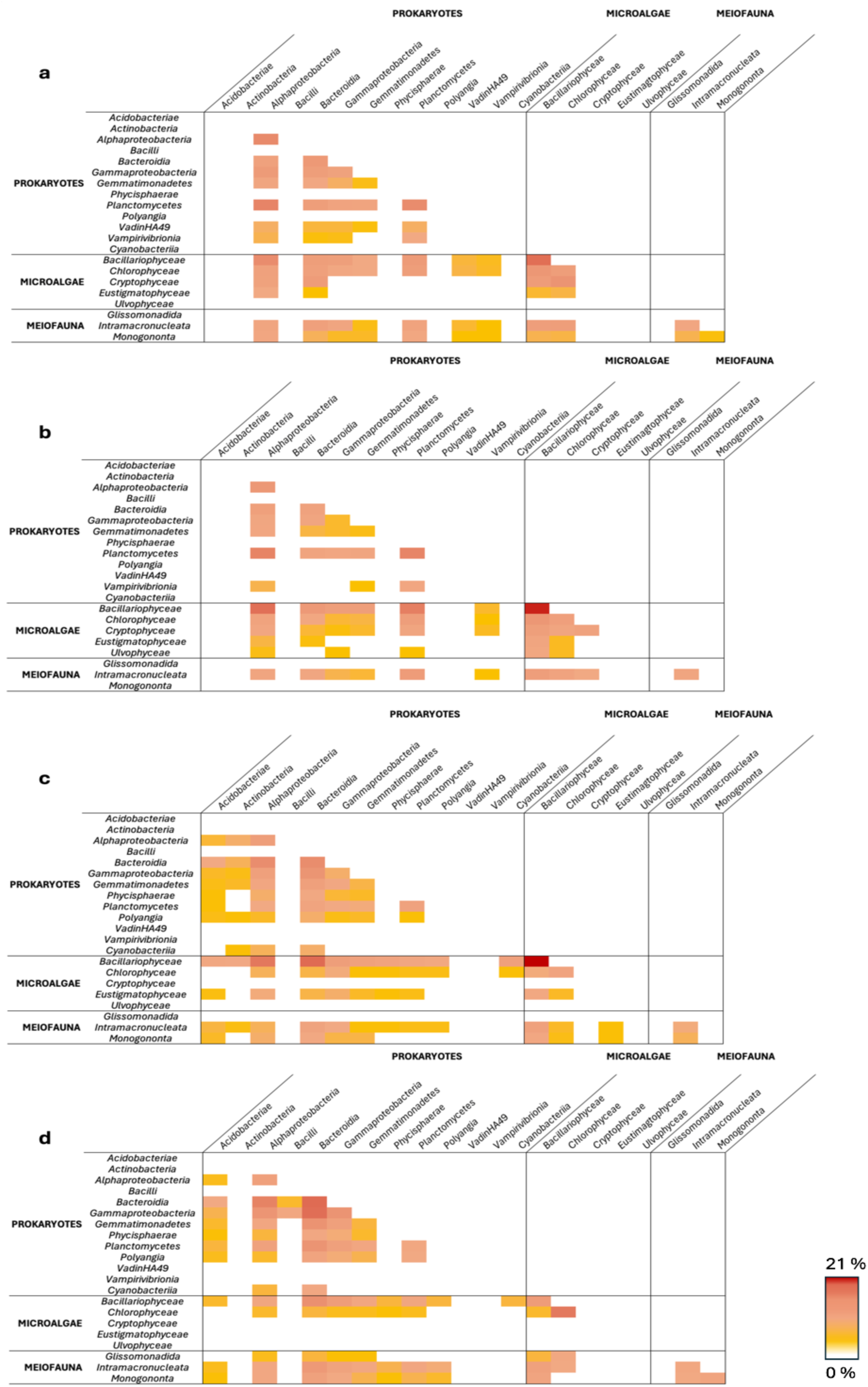
Matrix representing interactions between microbial taxa at the class taxonomical level. The gradient of color (from yellow (low proportions) to red (high proportions)) indicates the percentage of interactions attributed to a pair of taxa related to the total number of interactions. Each heatmap corresponds to a condition of Co exposure (Control (**a**), 0.1 µM Co (**b)**, 0.5 µM Co (**c**) and 1 µM Co (**d**)), including the biofilms at different stages of growth (from D7 to D28). Microbial taxa (at class taxonomical level) involved in less than 1% of the total interactions were removed for the analysis.

## 4. Discussion

As hotspots of microbial diversity, biofilms are functional consortia involved in nutrient and major biogeochemical cycles such as carbon, nitrogen, and phosphorus (3,4). All of these fluxes depend on the microbial interactions occurring locally. More importantly, interactions between autotrophic and heterotrophic components are of ecological and biogeochemical importance, allowing the maintenance of healthy biofilms, nutrient fluxes in the trophic web and the functionality of ecosystems.

One of our recent studies investigated the effects of Co on the prokaryotic component of freshwater biofilms (25). Both the prokaryotic community’s structure and composition were reported to be impacted when biofilms were grown in the presence of 0.5 and 1 µM Co. More precisely, *Cyanobacteria* and *Planctomycetes* were found to be sensitive while the *Bacteroidetes* appeared resistant. Also, high Co concentrations affected the putative functional potential of the prokaryotic component of biofilms. Indeed, the potential for non-photosynthetic carbon fixation by prokaryotes increased when the photosynthetic potential was significantly reduced due to the sensitivity of *Cyanobacteria* to Co. However, microalgae are key members of biofilms and can supply *Cyanobacteria* for phototrophic carbon fixation. Therefore, further investigation of the microeukaryotic component of these biofilms in order to provide a better understanding of the overall biofilm responses to Co was necessary.

Overall, the microeukaryotic community was sensitive to Co. It is worth noting that microeukaryotic and prokaryotic communities of the same biofilms were affected in their composition at similar Co concentrations. This corroborates the observation that microbial communities are spatially and functionally associated in biofilms and act as a single functional unit (40). Consequently, a contaminant can directly or indirectly impact microorganisms due to their close relationships. Interestingly, the *Bacillariophyta,* one of the dominating microalgae in the control biofilms, appeared particularly sensitive to Co. Along with the *Cyanobacteria,* which sensitivity has already been demonstrated (24,25), the phototrophic primary producers were the main metabolic group inhibited at 1 µM of Co. A canonical correspondence analysis (CCA) using previous published prokaryotic community diversity data further demonstrated the strong influence of phototrophic microorganisms on prokaryotic structure (Figure 5). The *Cyanobacteria* and *Bacillariophyceae* members were found to shape the prokaryotic communities in biofilms growing in control and 0.1 µM Co conditions with positive relationships. In contrast, the prokaryotic community structure of biofilms growing at 0.5 and 1 µM Co were influenced mainly by Co contamination but also by *Chlorophyceae* or *Cryptophyceae*.

**Figure 5:**
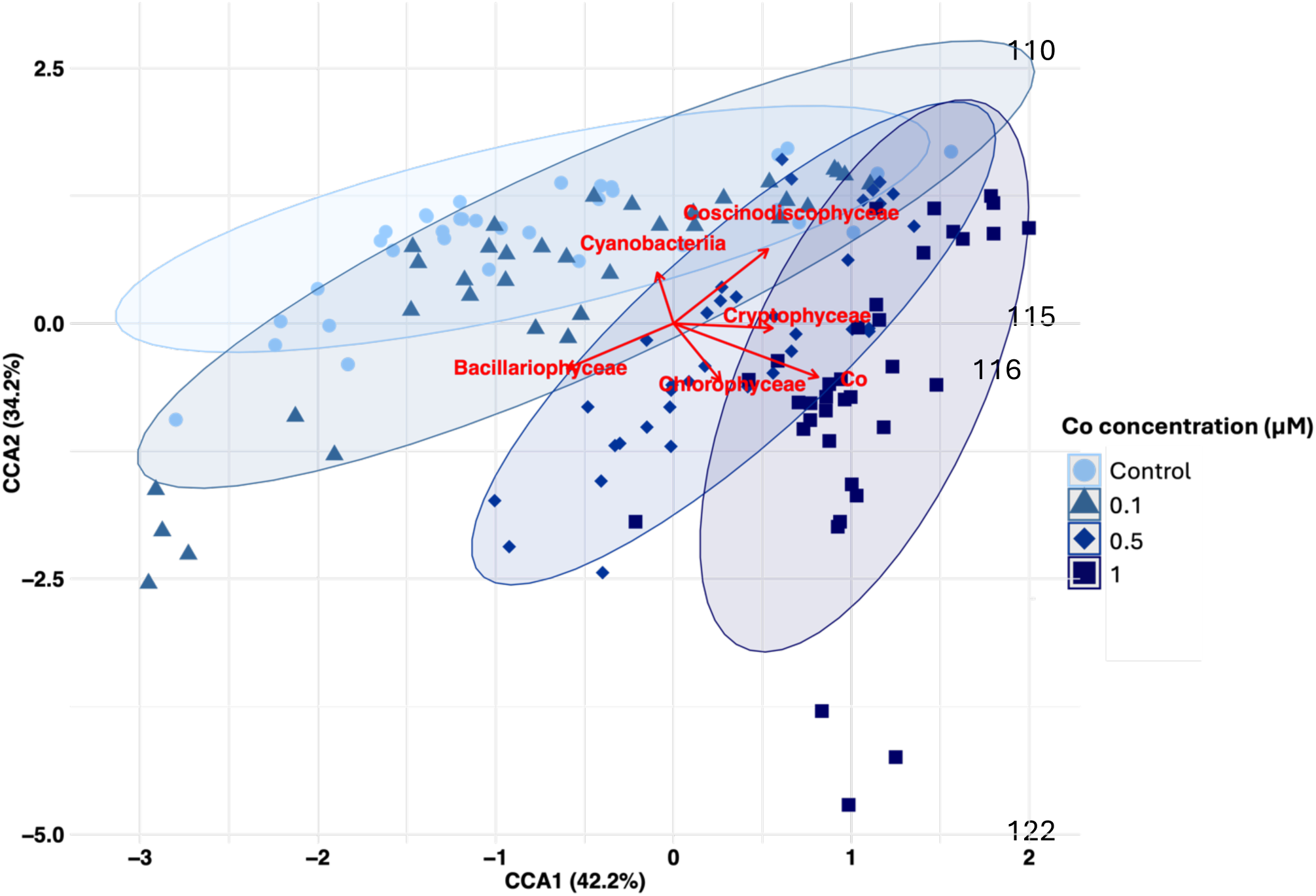
Biplot of canonical correspondence analysis (CCA) between prokaryotic community structure and dominant phototrophic taxa within biofilms growing at different Co exposure concentrations (control conditions with background Co concentration, 0.1, 0.5 or 1 µM Co). *Cyanobacteria, Bacillariophyceae, Chlorophyceae, Cryptophyceae, Coscinodiscophyceae* and Co concentrations were used as explanatory variables. The length and direction of the arrows indicate the strength and influence of the explanatory variables on the prokaryotic community structure of samples (colored depending on Co concentration, darker indicates higher Co concentration). No multicollinearity was observed between variables after tests using the Variance Inflation Factor (VIF). The ellipses represent the 95% confidence intervals for each Co concentration exposed samples. All explanatory variables represented are statistically significant (p < 0.05).

The co-occurrence network approach, which has drawn increasing attention over the last decade, provides an integrated view of all the potential interactions between organisms within a community. It also allows the identification of keystones, which are taxa that play crucial roles in maintaining the community, regardless of their abundance (41,42). Biofilms exposed to high Co contamination were characterized by networks with higher centralization around a few important nodes and a lower connectivity of microbial communities most likely due to the sensitivity of major primary producers. Such modifications of network topology were also observed in marine biofilms exposed to copper or high amounts of nutrients (43,44). In our study, network centralization and reduced connectivity may result from an altered autotroph-heterotroph coupling as suggested by microalgae sensitivity and the decrease of interactions involving microalgae. Along with our previous work where a significantly reduced photosynthetic potential was associated with an increasing potential for alternative carbon fixation pathways by prokaryotes (25), it appears that the Co modifies the potential metabolic functioning of biofilms by reducing photosynthesis. Thus, the heterotrophic community should adapt to the modification of the quality and quantity of produced organic matter as well as to Co presence.

Cobalt exposure led to smaller and more centralized networks, which are associated to low network complexity (45). However, the networks had higher density and cluster coefficient values at 1 µM Co, as reported in Cu-exposed biofilms (43) and soil microbial communities exposed to metals (9,46). High density means high number of connections per node (47) while an increasing cluster coefficient represents the predisposition to form sub-network and favor the interactions between species (48). We suggest that the increase of the number of connections would be a way for communities to favor resource access in the network under stress conditions ensuring support for biofilm growth and functionality. This hypothesis is supported by the decreasing network diameter and path length representing how quickly a community can respond to a stimulus (49) and how fast two nodes connect and pass information between them (50), respectively. Cobalt exposure also induced a shift in the network composition with an increasing prevalence of prokaryotes in the network structure and the microbial interactions while microalgae tended to be less numerous and less involved in interactions. The strong inhibition of photosynthetic microorganisms’ concomitant with shifts in the organization of the co-occurrence networks revealed that the inhibition of phototrophic primary producers in Co-exposed biofilms was probably the main cause of the modification of the heterotrophic community. Interestingly, the *Bacteroidetes* described as resistant under Co contamination, were drivers in the network interactions and part of the keystone taxa when the photoautotrophic function was altered at 1 µM Co. Indeed, members of the *Bacteroidota* phylum have already been described as dominant players in the degradation of suspended particles and refractory organic molecules present in their surroundings (1,51,52), making them available for other heterotrophs in response to photoautotrophic primary production impairment.

Keystone taxa are described as species that have a critical role in the structure and functioning of ecosystems irrespective of their abundance (34,53,54). They are essential for the stability of systems, and other organisms depend on their presence despite not being abundant when compared with other taxa in the community (55). In our study, we demonstrated that Co contamination led to a centralization of networks around a few keystones. A decreasing number of keystones was also described for soil microbial communities as a response to agricultural intensification (8). The shift from autotrophic-heterotrophic driven networks in natural biofilms to heterotrophic-dominated networks under high Co contamination was also supported by the affiliation of keystones in biofilms grown at 1 µM Co, which were 25% meiofauna and 75% prokaryotes including members of *Bacteroidetes, Gammaproteobacteria,* and *Gemmatimonadetes* (OTU25). These taxa exhibited resistance to Co (25) and, thanks to their metabolic abilities, contributed to the formation of new sub-networks when exposed to Co to favor the stability and functionality of the biofilms.

Interestingly, the OTU25 affiliated with *Gemmatimonadales* was the only OTU described as a keystone in both control conditions and at the highest Co concentration with a shift in the organization of its sub-network between the two conditions. *Gemmatimonadota*-affiliated members have been described in various environments, including aquatic habitats. Although the understanding of the metabolic properties of members of this phylum is poor with only 6 species described, cultured *Gemmatimonadota* are known as anoxygenic photoheterotrophs/heterotrophs, with a versatile metabolism (56–58). This metabolic versatility might allow the *Gemmatimonas* OTU25 to take advantage of the modification of organic matter production and availability (59,60). Indeed, as a heterotroph, OTU25 would use the organic carbon from its surroundings as carbon source and its phototrophic capabilities would also allow the use of light as an alternative energy source in order to compensate for the change in organic matter quality. Then, its resistance towards Co contamination would take part in the maintenance of biofilms in rivers, playing a pivotal role in the co-occurrence network.

Finally, as for the prokaryotic community, the microeukaryotic community composition remained different despite 35 days of recovery after stopping Co exposure (25). This absence of structural resilience in the microbial community composition suggests that the interactions between members of the biofilm will also remain affected in the long term and could lead to consequences up on the trophic chain and at the river scale. Reduced primary production productivity will not allow biofilms to cover long-term biotic needs and thereby reinforce the impairments on autotroph-heterotroph microbial interactions. In that dynamic, microbial composition will continue to be impacted and continue to favor the selection of competitive species affecting the biofilm structure and stability. The modification of EPS production (23) by microalgae or cyanobacteria is an example of disrupted functionality that can lead to a further reduction of biofilm quality and therefore the loss of its ecological function at the river scale including, nutrient cycling, water quality maintenance, and resources for higher trophic levels organisms.

In conclusion, this work demonstrated that the microbial connectivity and the abundance of keystones were higher in control biofilms indicating a great ecological stability. However, the presence of Co in rivers led to smaller networks dominated by prokaryotes. The adaptation of biofilms to Co exposure results from a cascade of effects in response to Co contamination which then impacts autotroph-heterotroph coupling.

## Acknowledgments

This research was funded by the Partnership Chair E2S-UPPA-TotalEnergies-Rio Tinto (ANR-16-IDEX-0002). We thank the members of TotalEnergies Environment & Sustainable Development Team at Pôle d’Études et de Recherches (PERL, Lacq, France) for the access to Pilot Rivers facilities and experimental assistance during the mesocosm experiment. Amplicon sequencing was performed by PGTB (Bordeaux, France) with the assistance of Zoé Delporte and Erwan Guichoux. We thank Claire Gassie (Université de Pau et des Pays de l’Adour, IPREM, Pau, France) for her technical assistance during laboratory work and characterization of microbial communities. We also thank François Rigal (Université de Pau et des Pays de l’Adour, IPREM, Pau, France) for its help in statistical analyses and Olivier Pringault (Institut Méditerranéen d’Océanologie, Marseille, France) for his critical reading on the earlier version of the manuscript.

## Data Availability Statement

Raw DNA sequences were submitted to the National Center for Biotechnology Information as a Sequence Read Archive (SRA) under the accession number PRJNA1166897 and will be available on 2025-01-01.

## Associated content

**Supporting Information:** Includes supporting figures (Figures S1-S7) and supporting table (Table S1)

**Appendix A**: Additional tables (Table A1-Table A11)

