## Supplementary Material for "Cobalt induces the set-up of new structural networks in river biofilms: Impairment of autotrophic-heterotrophic coupling"

Postal address: Technopôle Hélioparc, 2 Av. Du Président Pierre Angot, 64053 Pau Cedex 9

This **supplementary information file** includes:

- Supplementary figures (Figures S1-S7)
- Supplementary table (Table S1)
- Titles of additional tables available in **Appendix A** (Table A1- Table A11).

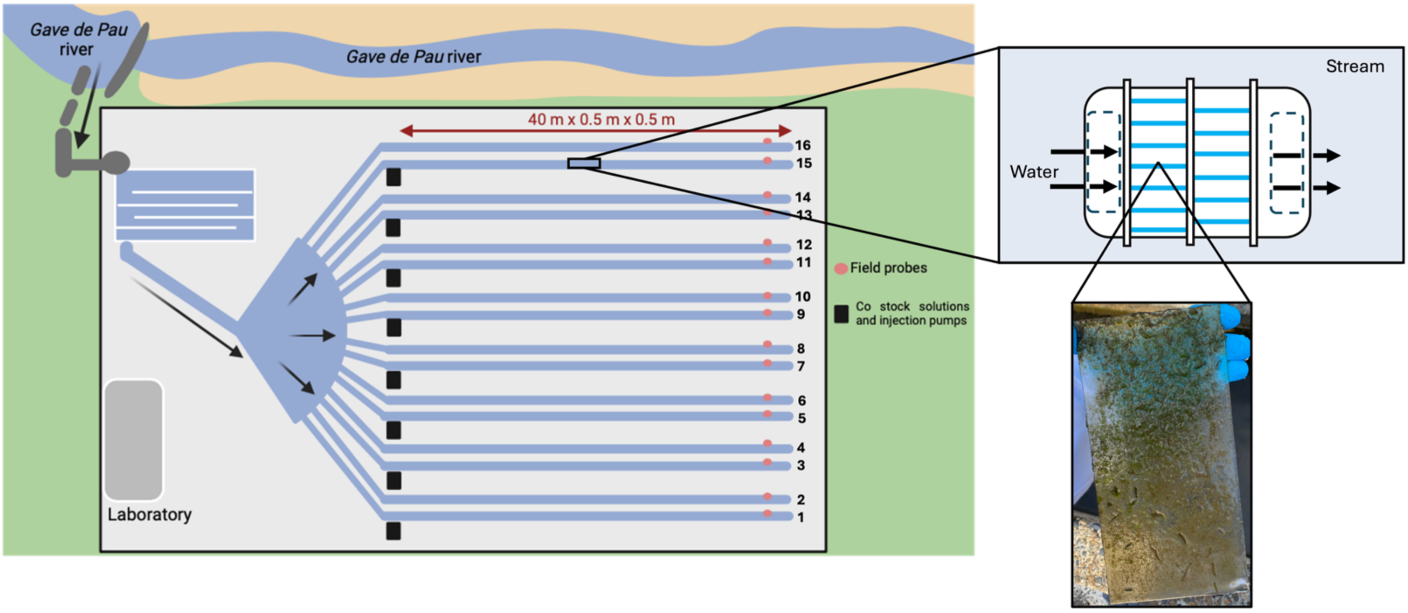


**Figure S1**: The Pilot Rivers facility of TotalEnergies is outdoor artificial streams located in Lacq, France (1). The Pilot Rivers consist of 16 artificial streams (40 m long, 0.5 m wide, 0.5 m deep and with a flow rate set at 7.5 $m^3.h^{-1}$ for the experimentation) fed continuously by the *Gave de Pau River*. Four treatments of Co contamination were continuously applied in triplicates of streams after random selection: streams 6, 8 and 14 for control conditions with background Co concentrations of the *Gave de Pau River;* channels 5, 10 and 15 for 0.1 µM Co; channels 2, 3 and 9 for 0.5 µM Co and channels 7, 11 and 12 for 1 µM Co. Solutions of cobalt (Cobalt(II) chloride hexahydrate, 98%, Thermo Scientific Chemicals) were injected continuously (71 $mL.h^{-1}$) in streams with volumetric (membrane) pumps (Prominent), for 28 days of exposure. At each end of stream, field probes (Hach, IA, USA) were used to measure the temperature (°C), pH, conductivity, oxygen saturation and dissolved oxygen before sampling. A plastic box containing 13 sterile glass slides for biofilm colonization was deposited parallel to the flow in each stream at a distance of 15 meters from the injection pumps. The water at the outlet of the streams was directed into the phytotreatment lagoon (reedbeds) to reduce the Co concentration during the entire experiment. Downstream the phytotreatment, the water was sampled and analysed for anomalies before being discharged.

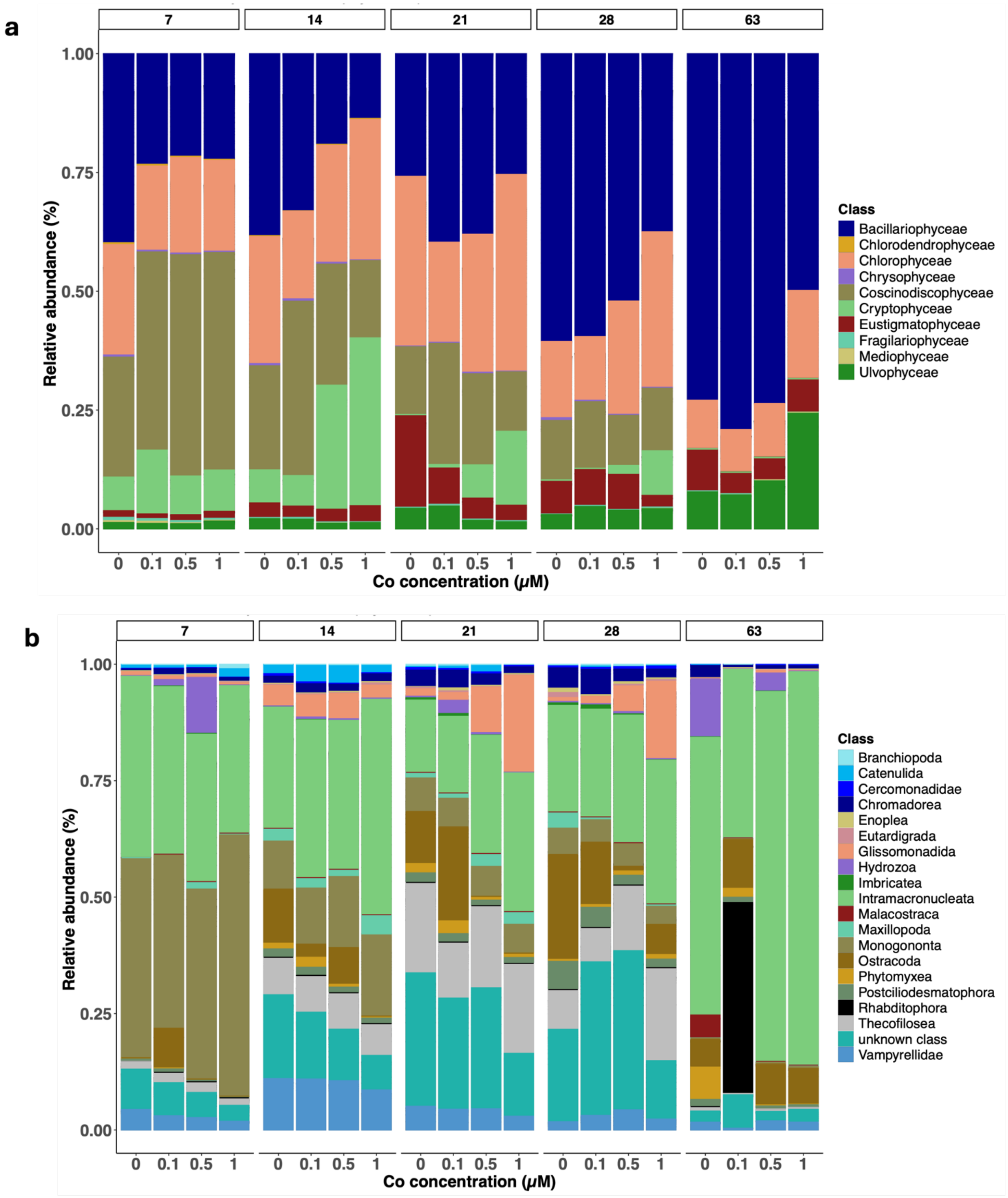


**Figure S2:** Microeukaryotic community diversity of microalgae (**a**) and meiofauna (**b**) during biofilm colonization (D7, D14, D21, D28 and D63-R) as a function of Co concentration (control, 0.1 µM, 0.5 µM and 1 µM)

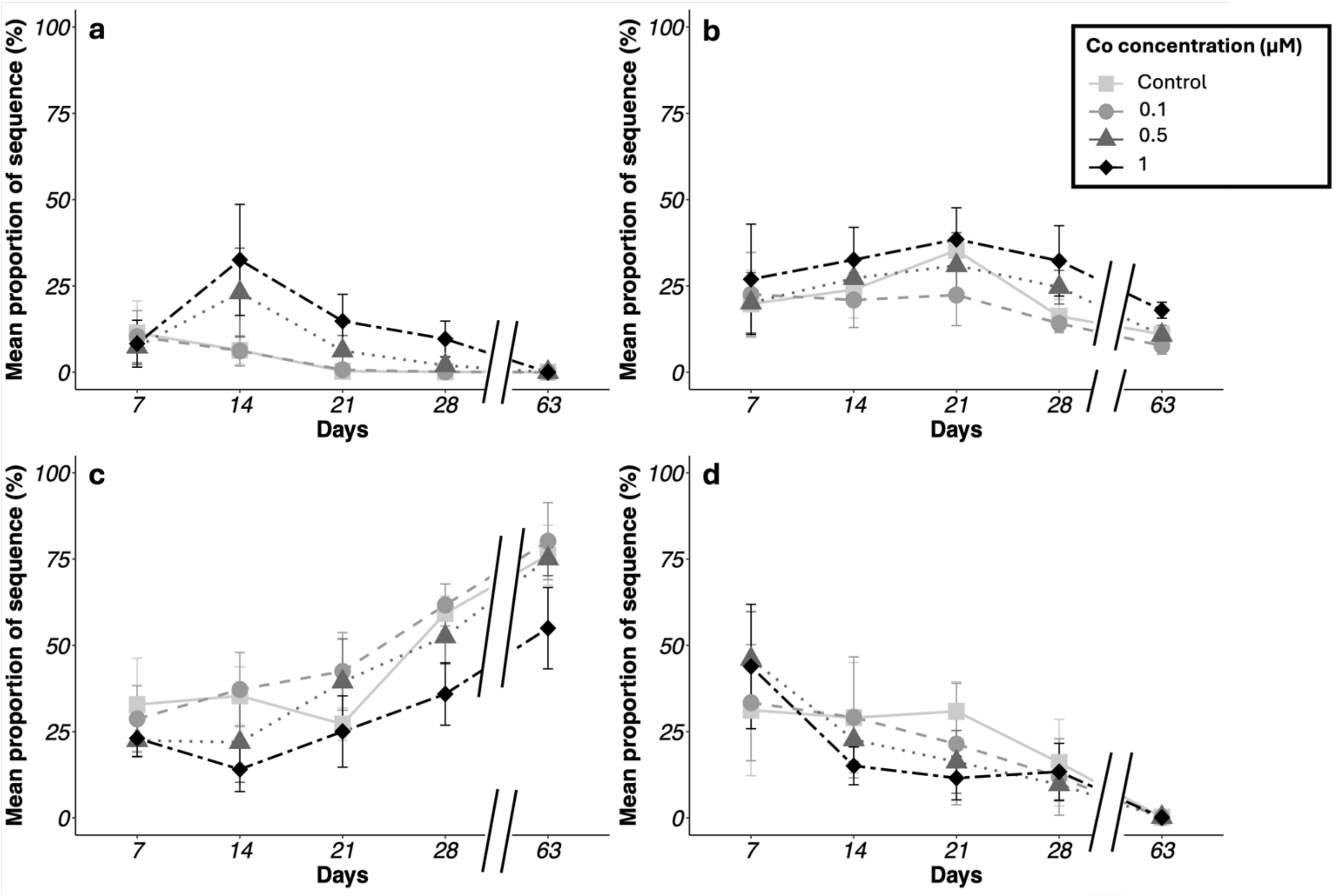


**Figure S3:** Relative proportions of *Cryptophyceae* (**a**), *Chlorophyceae* (**b**)*, Bacillariophycea* (**c**) and *Coscinodiscophyceae* (d) classes of microalgal community within biofilms as a function of exposure time at several Co concentrations. The shape of points and lines indicates the increasing Co concentrations by comparison with control conditions (■: Control conditions; ●: 0.1 µM Co; ▲: 0.5 µM Co; ◆: 1 µM Co).

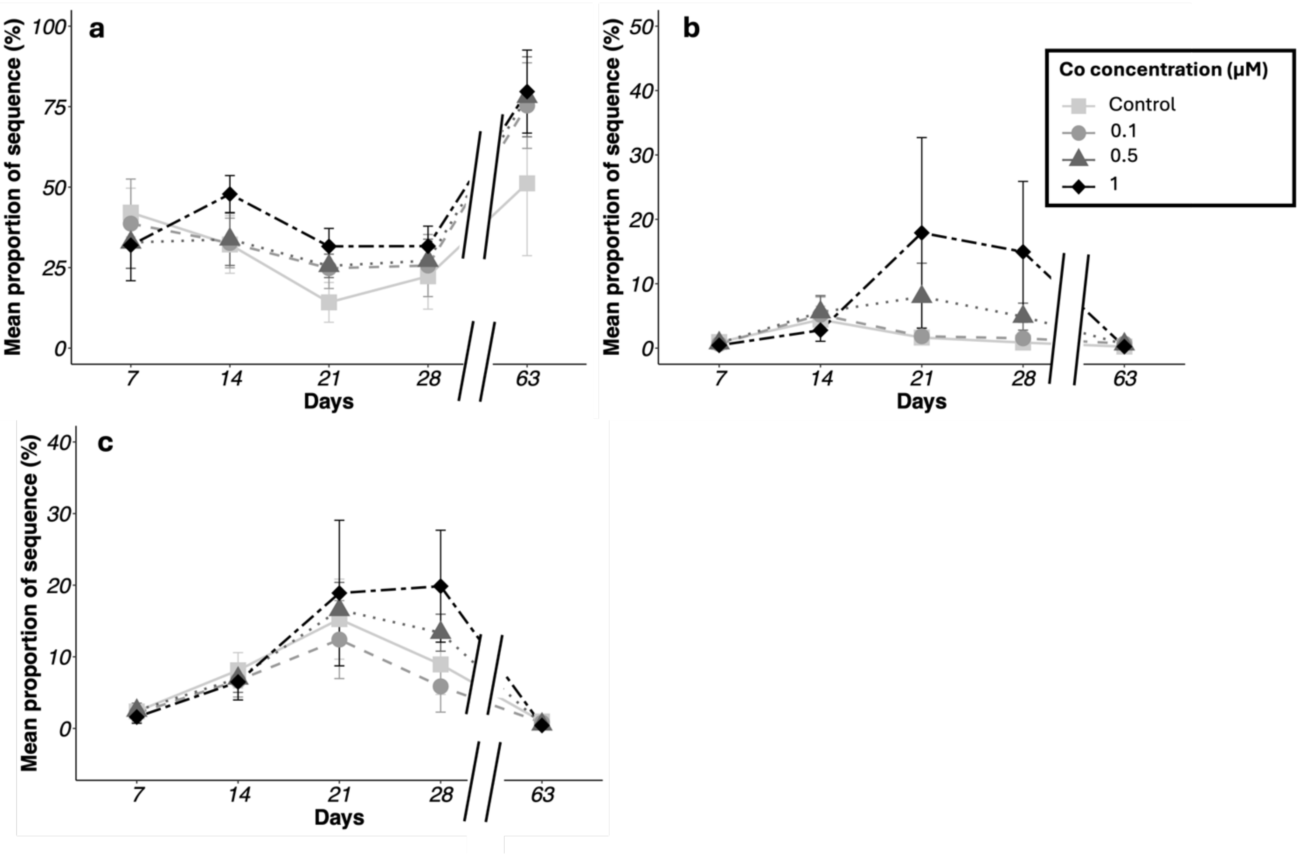


**Figure S4:** Relative proportions of *Intramacronucleata* (**a**), *Glassimonadida* (**b**), and *Thecofilosea* (**c**) classes of meiofaunal community within biofilms as a function of exposure time at several Co concentrations.. The shape of points and lines indicates the increasing Co concentrations by comparison with control conditions (■: Control conditions; ●: 0.1 µM Co; ▲: 0.5 µM Co; ◆: 1 µM Co).

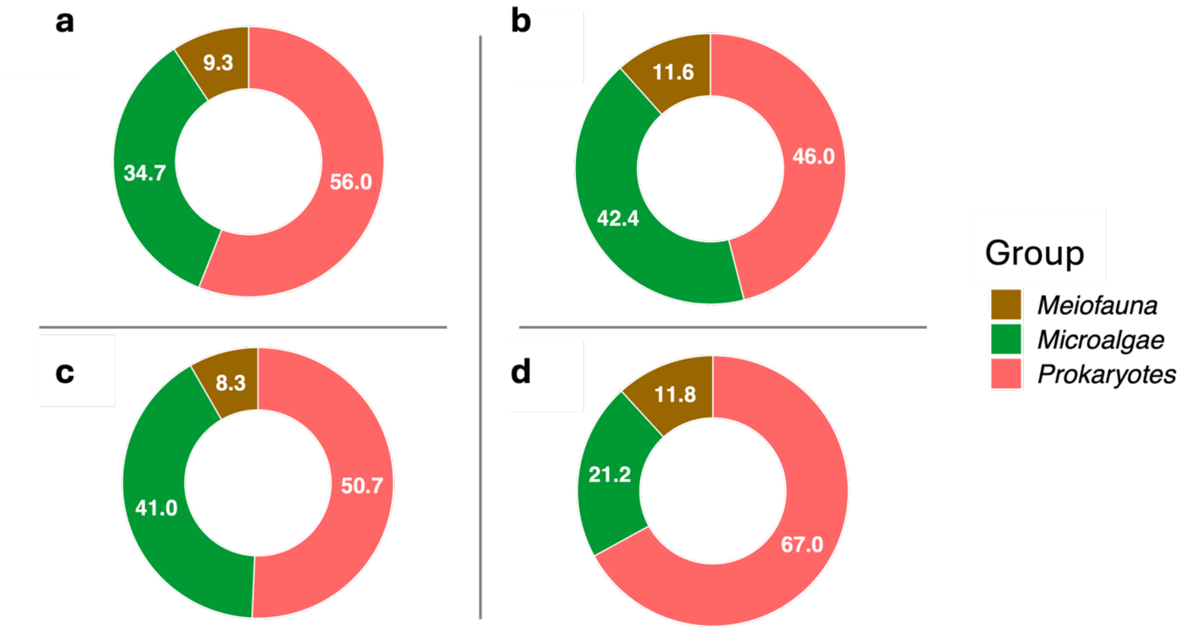


**Figure S5:** Proportions of nodes associated with studied microbial groups (prokaryotes, microalgae and meiofauna) as a function of exposure conditions for biofilms grown in conditions of control (**a**), 0.1 µM Co (**b**), 0.5 µM Co (**c**) and 1 µM Co (**d**).

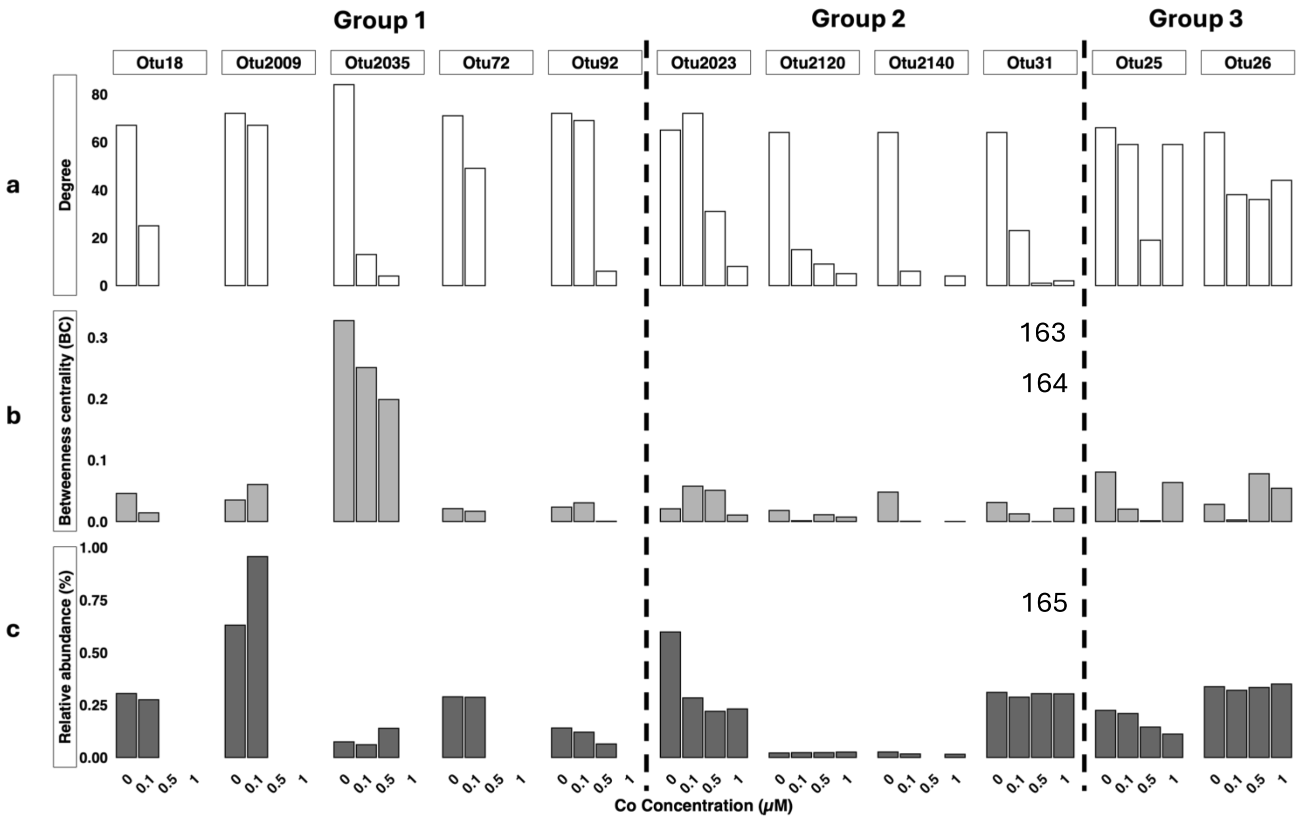


**Figure S6:** Degree (**a**) and betweenness centrality (**b**) parameters of the keystone OTUs identified in the control conditions in the networks produced by biofilms grown in the presence of 0.1 μM Co, 0.5 μM Co and 1 μM Co. Relative abundance (**c**) of control keystone OTUs in the global microbial community as a function of Co exposure concentrations.

**a**

| | | + | - |
|---|---|---|---|
| Prokaryotes | ***Alphaproteobacteria*** | **7** | **1** |
| | *Caulobacterales* | 1 | 0 |
| | *Rhizobiales* | 2 | 0 |
| | *Rhodobacterales* | 3 | 0 |
| | *Rickettsiales* | 1 | 1 |
| | ***Bacteroidia*** | **0** | **3** |
| | *Flavobacteriales* | 0 | 2 |
| | *Sphingobacteriales* | 0 | 1 |
| | ***Gammaproteobacteria*** | **7** | **0** |
| | *Burkholderiales* | 1 | 0 |
| | *Gammaproteobacteria Incertae Sedis* | 1 | 0 |
| | *Xanthomonadales* | 5 | 0 |
| | ***Planctomycetes*** | **4** | **0** |
| | *Gemmatales* | 3 | 0 |
| | *Pirellulales* | 1 | 0 |
| | ***vadinHA49*** | **0** | **1** |
| | *unknown order* | 0 | 1 |
| Microalgae | ***Bacillariophyceae*** | **2** | **0** |
| | *unknown order* | 2 | 0 |
| | ***Chlorophyceae*** | **0** | **3** |
| | *Chlamydomonadales* | 0 | 2 |
| | *unknown order* | 0 | 1 |
| | ***Cryptophyceae*** | **0** | **3** |
| | *Cryptomonadales* | 0 | 3 |
| Meiofauna | ***Monogononta*** | **0** | **1** |
| | *Ploimida* | 0 | 1 |
| | ***Vampyrellidae*** | **0** | **1** |
| | *unknown order* | 0 | 1 |
| | **Total** | **20** | **13** |

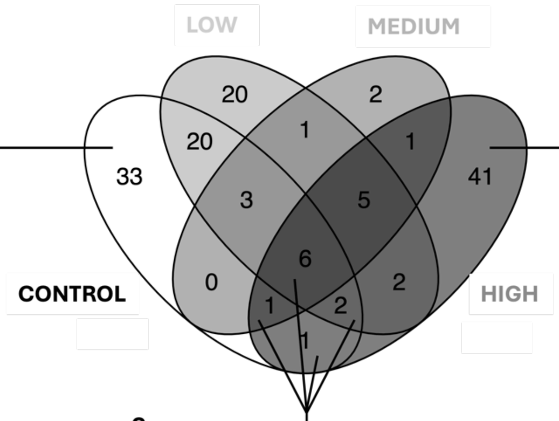


**b**

| | | + | - |
|---|---|---|---|
| Prokaryotes | ***Alphaproteobacteria*** | **1** | **5** |
| | *Rhodobacterales* | 0 | 3 |
| | *Rickettsiales* | 1 | 2 |
| | ***Bacteroidia*** | **10** | **2** |
| | *Chitinophagales* | 1 | 1 |
| | *Cytophagales* | 1 | 1 |
| | *Flavobacteriales* | 2 | 0 |
| | *Sphingobacteriales* | 6 | 0 |
| | ***Gammaproteobacteria*** | **4** | **3** |
| | *Burkholderiales* | 2 | 3 |
| | *Legionellales* | 1 | 0 |
| | *Xanthomonadales* | 1 | 0 |
| | ***Gemmatimonadetes*** | **1** | **0** |
| | *Gemmatimonadales* | 1 | 0 |
| | ***Planctomycetes*** | **3** | **0** |
| | *Pirellulales* | 3 | 0 |
| | ***Verrucomicrobiae*** | **1** | **0** |
| | *Opitutales* | 1 | 0 |
| Microalgae | ***Bacillariophyceae*** | **1** | **4** |
| | *unknown order* | 1 | 4 |
| | ***Chlorophyceae*** | **0** | **1** |
| | *Chlamydomonadales* | 0 | 1 |
| Meiofauna | ***Glissomonadida*** | **1** | **0** |
| | *unknown order* | 1 | 0 |
| | ***Intramacronucleata*** | **0** | **4** |
| | *Conthreep* | 0 | 2 |
| | *Litostomatea* | 0 | 1 |
| | *Spirotrichea* | 0 | 1 |
| | **Total** | **22** | **19** |

**c**

| | | + | - |
|---|---|---|---|
| Prokrayotes | ***Bacteroidia*** | **0** | **4** |
| | *Chitinophagales* | 0 | 4 |
| | ***Gemmatimonadetes*** | **2** | **0** |
| | *Gemmatimonadales* | 2 | 0 |
| | ***Planctomycetes*** | **1** | **0** |
| | *Gemmatales* | 1 | 0 |
| Microalgae | ***Cryptophyceae*** | **0** | **1** |
| | *Cryptomonadales* | 0 | 1 |
| Meiofauna | ***Intramacronucleata*** | **0** | **1** |
| | *Conthreep* | 0 | 1 |
| | ***Monogononta*** | **0** | **1** |
| | *Ploimida* | 0 | 1 |
| | **Total** | **3** | **7** |

**Figure S7:** Venn diagram of the number of interactions for the control and 1 µM Co keystone OTU25 as a function of Co exposure concentration. The type (positive (+)/ negative (-)) and taxonomy of the nodes connected to the OTU25 are presented in the tables for the interactions occurring in control conditions (**a**), highest exposure conditions (**b**), and for the interactions shared between control and high exposure conditions (**c**). Exposure conditions are named as follows: background Co concentration in the Gave de Pau River - Control; 0.1 µM Co - Low; 0.5 µM Co - Medium and 1 µM Co - High.

**Table S1**: List of the direct neighbors of the OTU25 in networks as a function of Co concentration. Class and orders are described, such as the number and type of interactions (positive or negative). The keystone character is represented by an underlined Co concentration.

| | Control | | 0.1 | | 0.5 | | 1 | |
|---|---|---|---|---|---|---|---|---|
| | + | - | + | - | + | - | + | - |
| • Acidobacteria | 0 | 0 | 0 | 0 | 0 | 0 | 1 | 0 |
| Bryobacterales | 0 | 0 | 0 | 0 | 0 | 0 | 1 | 0 |
| •Alphaprotrobacteria | 12 | 1 | 7 | 1 | 1 | 1 | 1 | 6 |
| Caulobacter | 1 | 0 | 0 | 0 | 0 | 0 | 0 | 0 |
| Rhizobiales | 2 | 0 | 0 | 0 | 0 | 0 | 0 | 0 |
| Rhodobacterales | 3 | 0 | 0 | 0 | 0 | 0 | 0 | 3 |
| Rickettsiales | 3 | 1 | 3 | 1 | 1 | 1 | 1 | 3 |
| Sphingomonadales | 3 | 0 | 4 | 0 | 0 | 0 | 0 | 0 |
| •Bacillariophyceae | 5 | 2 | 15 | 3 | 0 | 2 | 1 | 6 |
| Unknown order | 5 | 2 | 15 | 3 | 0 | 2 | 1 | 6 |
| •Bacteroidia | 0 | 7 | 0 | 5 | 1 | 3 | 10 | 6 |
| Chitinophagales | 0 | 4 | 0 | 5 | 0 | 3 | 0 | 5 |
| Cytophagales | 0 | 0 | 0 | 0 | 0 | 0 | 1 | 1 |
| Flavobacteriales | 0 | 2 | 0 | 0 | 1 | 0 | 3 | 0 |
| Sphingobacteriales | 0 | 1 | 0 | 0 | 0 | 0 | 6 | 0 |
| •Chlorophyceae | 0 | 6 | 0 | 3 | 0 | 0 | 0 | 1 |
| Chlamydomonadales | 0 | 5 | 0 | 3 | 0 | 0 | 0 | 1 |
| Unknown order | 0 | 1 | 0 | 0 | 0 | 0 | 0 | 0 |
| •Cryptophyceae | 0 | 5 | 0 | 1 | 0 | 0 | 0 | 1 |
| Cryptomonadales | 0 | 5 | 0 | 2 | 0 | 0 | 0 | 1 |
| •Eustigmatophyceae | 0 | 0 | 0 | 0 | 1 | 0 | 0 | 0 |
| Eustigmatales | 0 | 0 | 0 | 0 | 1 | 0 | 0 | 0 |
| •Gammaproteobacteria | 7 | 0 | 2 | 1 | 1 | 1 | 5 | 4 |
| Burkholderiales | 1 | 0 | 1 | 1 | 1 | 1 | 3 | 4 |
| Gammaproteobacteria Incertae Sedis | 1 | 0 | 0 | 0 | 0 | 0 | 0 | 0 |
| Legionellales | 0 | 0 | 0 | 0 | 0 | 0 | 1 | 0 |
| Pseudomonadales | 0 | 0 | 0 | 0 | 0 | 0 | 0 | 0 |
| Xanthomonadales | 5 | 0 | 1 | 0 | 0 | 0 | 1 | 0 |
| •Gemmatimonadetes | 2 | 0 | 2 | 0 | 2 | 0 | 3 | 0 |
| Gemmatimonadales | 2 | 0 | 2 | 0 | 2 | 0 | 3 | 0 |
| •Intramacronucleata | 0 | 1 | 0 | 5 | 0 | 0 | 0 | 6 |

| | | | | | | | | |
|---|---|---|---|---|---|---|---|---|
| Conthreep | 0 | 1 | 0 | 3 | 0 | 0 | 0 | 3 |
| Litostomatea | 0 | 0 | 0 | 2 | 0 | 0 | 0 | 2 |
| Spirotrichea | 0 | 0 | 0 | 0 | 0 | 0 | 0 | 1 |
| •Monogononta | 0 | 3 | 0 | 2 | 0 | 2 | 0 | 1 |
| Ploimida | 0 | 1 | 0 | 2 | 0 | 2 | 0 | 1 |
| •Phycisphaerae | 0 | 0 | 1 | 0 | 1 | 0 | 1 | 0 |
| Tepidisphaerales | 0 | 0 | 1 | 0 | 1 | 0 | 1 | 0 |
| •Planctomycetes | 13 | 0 | 11 | 0 | 3 | 0 | 4 | 0 |
| Gemmatales | 5 | 0 | 2 | 0 | 1 | 0 | 1 | 0 |
| Isosphaerales | 1 | 0 | 0 | 0 | 0 | 0 | 0 | 0 |
| Pirellulales | 6 | 0 | 6 | 0 | 2 | 0 | 3 | 0 |
| Planctomycetales | 2 | 0 | 3 | 0 | 0 | 0 | 0 | 0 |
| •VadinHA49 | 0 | 1 | 0 | 0 | 0 | 0 | 0 | 0 |
| Unknown order | 0 | 1 | 0 | 0 | 0 | 0 | 0 | 0 |
| •Vampyrellidae | 0 | 1 | 0 | 0 | 0 | 0 | 0 | 0 |
| Unknown order | 0 | 1 | 0 | 0 | 0 | 0 | 0 | 0 |
| •Verrucomicrobiae | 0 | 0 | 0 | 0 | 0 | 0 | 1 | 0 |
| Opitutales | 0 | 0 | 0 | 0 | 0 | 0 | 1 | 0 |
| Total | 39 | 27 | 38 | 21 | 10 | 9 | 28 | 31 |
| Total | 66 | | 59 | | 19 | | 59 | |

**References supplementary information:**


(1) Cailleaud, K., Bassères, A., Gelber, C., Postma, J.F., Ter Schure, A.T.M., Leonards, P.E.G., Redman, A.D., Whale, G.F., Spence, M.J., Hjort, M., 2019. Investigating predictive tools for refinery effluent hazard assessment using stream mesocosms. Enviro Toxic and Chemistry 38, 650–659. https://doi.org/10.1002/etc.4338

Additional tables are available in the **Appendix A** file (Table A1-Table A11).

**Table A1:** 18S rRNA gene copy numbers at different stages (days) of biofilm colonization at different Co concentrations.

**Table A2:** Percentage of major groups within eukaryotic community in biofilms exposed to different Co concentrations as a function of the colonization stage (days)

**Table A3:** Alpha-diversity indexes of microalgal community within growing biofilms colonized with different Co concentrations and as a function of the colonization stage (days)

**Table A4:** Indexes of alpha-diversity of meiofauna within growing biofilms colonized in presence of Co and as a function of the colonization stage (days)

**Table A5:** Cobalt effect on beta-diversity of microalgal community over biofilms colonization at different Co concentrations. Groupe comparison scores and significances levels defined after Pairwise permanova test. Sampling times and Co concentration are annoted as follows: T1 (Day 7), T2 (Day14), T3 (Day21), T4 (Day28), T5 (Recovery), C1 (Control), C2 (0.1μM Co), C3 (0.5 μM Co), C4 (1 μM Co). Significances of tests are represented as follows: . ($p< 0.05$), * ($p<0.01$) and ** ($p<0.001$).

**Table A6:** Cobalt effect on beta-diversity of meiofaunal community over biofilms colonization at different Co concentrations. Groupe comparison scores and significances levels defined after Pairwise permanova test. Sampling times and Co concentration are annoted as follows: T1 (Day 7), T2 (Day14), T3 (Day21), T4 (Day28), T5 (Recovery), C1 (Control), C2 (0.1μM Co), C3 (0.5 μM Co), C4 (1 μM Co). Significances of tests are represented as follows: . ($p< 0.05$), * ($p<0.01$) and ** ($p<0.001$).

**Table A7:** Keystone OTUs identified in control biofilms and their relative abundance depending on Co exposure conditions.

**Table A8:** Percentage of interactions of the main taxa in the control network. Values indicate the proportion of interactions in which taxa are involved at least once, considering that the interaction can be either single-taxum or bi-taxa. The network was constructed including the communities from D7 to D28. Microbial taxa (at class taxonomical level) involved in less than 1% of the total interactions were removed from the analysis.

**Table A9:** Percentage of interactions of the main taxa in the network from 0.1 μM Co condition. Values indicate the proportion of interactions in which taxa are involved at least once, considering that the interaction can be either single-taxum or bi-taxa. The network was constructed including the communities from D7 to D28. Microbial taxa (at class taxonomical level) involved in less than 1% of the total interactions were removed from the analysis.

**Table A10:** Percentage of interactions of the main taxa in the network from 0.5 μM Co condition. Values indicate the proportion of interactions in which taxa are involved at least once, considering that the interaction can be either single-taxum or bi-taxa. The network was constructed including the communities from D7 to D28. Microbial taxa (at class taxonomical level) involved in less than 1% of the total interactions were removed from the analysis.

**Table A11:** Percentage of interactions of the main taxa in the network from 1 μM Co condition. Values indicate the proportion of interactions in which taxa are involved at least once, considering that the interaction can be either single-taxum or bi-taxa. The network was constructed including the communities from D7 to D28. Microbial taxa (at class taxonomical level) involved in less than 1% of the total interactions were removed from the analysis.
